# FLT3-ITD transduces autonomous growth signals during its biosynthetic trafficking in acute myelogenous leukemia cells

**DOI:** 10.1101/2021.01.01.424454

**Authors:** Kouhei Yamawaki, Isamu Shiina, Takatsugu Murata, Satoru Tateyama, Yutarou Maekawa, Mariko Niwa, Motoyuki Shimonaka, Koji Okamoto, Toshihiro Suzuki, Toshirou Nishida, Ryo Abe, Yuuki Obata

**Author notes:** **Corresponding author**: Yuuki Obata, Ph.D., Division of Cancer Differentiation, National Cancer Center Research Institute, Tsukiji, 5-1-1, Chuo-ku, Tokyo,104-0045,Japan, **Co-corresponding author** Ryo Abe, M.D., Ph.D., SIRC, Teikyo University, Kaga, 2-11-1, Itabashi-ku, Tokyo, 173-8605, Japan. **Grant Support** Japan Society for the Promotion of Science (18K07208 to YO, 19H03722 to TN and 20K08719 to RA) Friends of Leukemia Research Fund (to YO) Kawano Masanori Memorial Public Interest Incorporated Foundation for Promotion of Pediatrics (to YO) Ichiro Kanehara Foundation for the Promotion of Medical Sciences and Medical Care (To YO). **E-mail contacts**, Kouhei Yamawaki, Isamu Shiina, Takatsugu Murata, Satoru Tateyama, Yutarou Maekawa, Mariko Niwa, Motoyuki Shimonaka, Koji Okamoto, Toshihiro Suzuki, Toshirou Nishida.

## Abstract

FMS-like tyrosine kinase 3 (FLT3) in hematopoietic cells binds to its ligand at the plasma membrane (PM), then transduces growth signals. *FLT3* gene alterations that lead the kinase to assume its permanently active form, such as *internal tandem duplication (ITD)* and *D835Y* substitution, are found in 30~40% of acute myelogenous leukemia (AML) patients. Thus, the drugs for molecular targeting of FLT3 mutants have been developed for the treatment of AML. Several groups have reported that compared with wild-type FLT3 (FLT3-wt), FLT3 mutants are retained in organelles, resulting in low levels of PM localization of the receptor. However, the precise subcellular localization of mutant FLT3 remains unclear, and the relationship between oncogenic signaling and the mislocalization is not completely understood. In this study, we show that in cell lines established from AML patients, endogenous FLT3-ITD but not FLT3-wt clearly accumulates in the perinuclear region. Our co-immunofluorescence assays demonstrate that Golgi markers are co-localized with the perinuclear region, indicating that FLT3-ITD mainly localizes to the Golgi region in AML cells. FLT3-ITD biosynthetically traffics to the Golgi apparatus and remains there in a manner dependent on its tyrosine kinase activity. A tyrosine kinase inhibitor midostaurin (PKC412) markedly decreases in FLT3-ITD retention and increases in the PM levels of the mutant. FLT3-ITD activates downstream in the endoplasmic reticulum (ER) and the Golgi apparatus during its biosynthetic trafficking. Results of our trafficking inhibitor treatment assays show that FLT3-ITD in the ER activates STAT5, whereas that in the Golgi can cause the activation of AKT and ERK. We provide evidence that FLT3-ITD signals from the early secretory compartments before reaching the PM in AML cells.

## Introduction

FLT3 is a member of the type III receptor type tyrosine kinase (RTK) family and is expressed in the PM of hematopoietic cells (Lemmon & Schlessinger, 2010; Tsapogas et al., 2017; Kazi & Rönnstrand, 2019). Upon stimulation with FLT3 ligand, the receptor undergoes dimerization and autophosphorylates its tyrosine residues, such as Tyr591 and Tyr842 (Choudhary et al., 2009; Köthe et al., 2013; Kazi & Rönnstrand, 2019). Subsequently, it activates downstream molecules, such as AKT, extracellular signal-regulated kinase (ERK), and signal transducers and activators of transcription (STAT) proteins (Meshinchi & Appelbaum, 2009; Kazi & Rönnstrand, 2019). Activation of these cascades results in the growth and differentiation of host cells, leading to normal hematopoiesis (Tsapogas et al., 2017). Therefore, gain-of-function mutations of *FLT3* cause autonomous proliferation of myeloid cells, resulting in the development of AML (Daver et al., 2019; Kiyoi et al., 2020).

FLT3 is composed of an N-terminal extracellular domain, a transmembrane region, a juxta-membrane (JM) domain, and a C-terminal cytoplasmic tyrosine kinase domain (Lemmon & Schlessinger, 2010; Kazi & Rönnstrand, 2019; see Figure 1A). Alterations of the *FLT3* gene that lead the kinase to constitutive activation are seen in 30~40% of AML cases (Meshinchi & Appelbaum, 2009; Kiyoi et al., 2020). Internal tandem duplication (ITD) into the JM region of FLT3 interferes with its auto-inhibitory ability (Kiyoi et al., 2002). In addition, a D835Y substitution in the FLT3 activation loop stabilizes the tyrosine kinase domain in an active state (Yamamoto et al., 2001; Lemmon & Schlessinger, 2010). Thus, signal transduction pathways from FLT3 mutants have been investigated (Meshinchi & Appelbaum, 2009; Chatterjee et al., 2015; Chougule et al., 2016; Heydt et al., 2018), and molecular targeting drugs for blocking the mutants have been developed for the treatment of AML patients (Gallogly et al., 2017; Daver et al., 2019; Kiyoi et al., 2020). Previous studies showed that FLT3-ITD accumulates in the wrong compartments, resulting in low amounts of the mutant in the PM, compared with the allocation of FLT3-wt (Schmidt-Arras et al., 2005; Koch et al., 2008; Choudhary et al., 2009; Köthe et al., 2013; Kellner et al., 2020). Although FLT3-ITD is suggested to activate STAT5 soon after synthesis (Schmidt-Arras et al., 2009; Choudhary et al., 2009; Tsitsipatis et al., 2017; Takahashi, 2019), the precise subcellular localization of the mutant and the relationship between the mislocalization and growth signals remain unclear.

**Figure 1.**
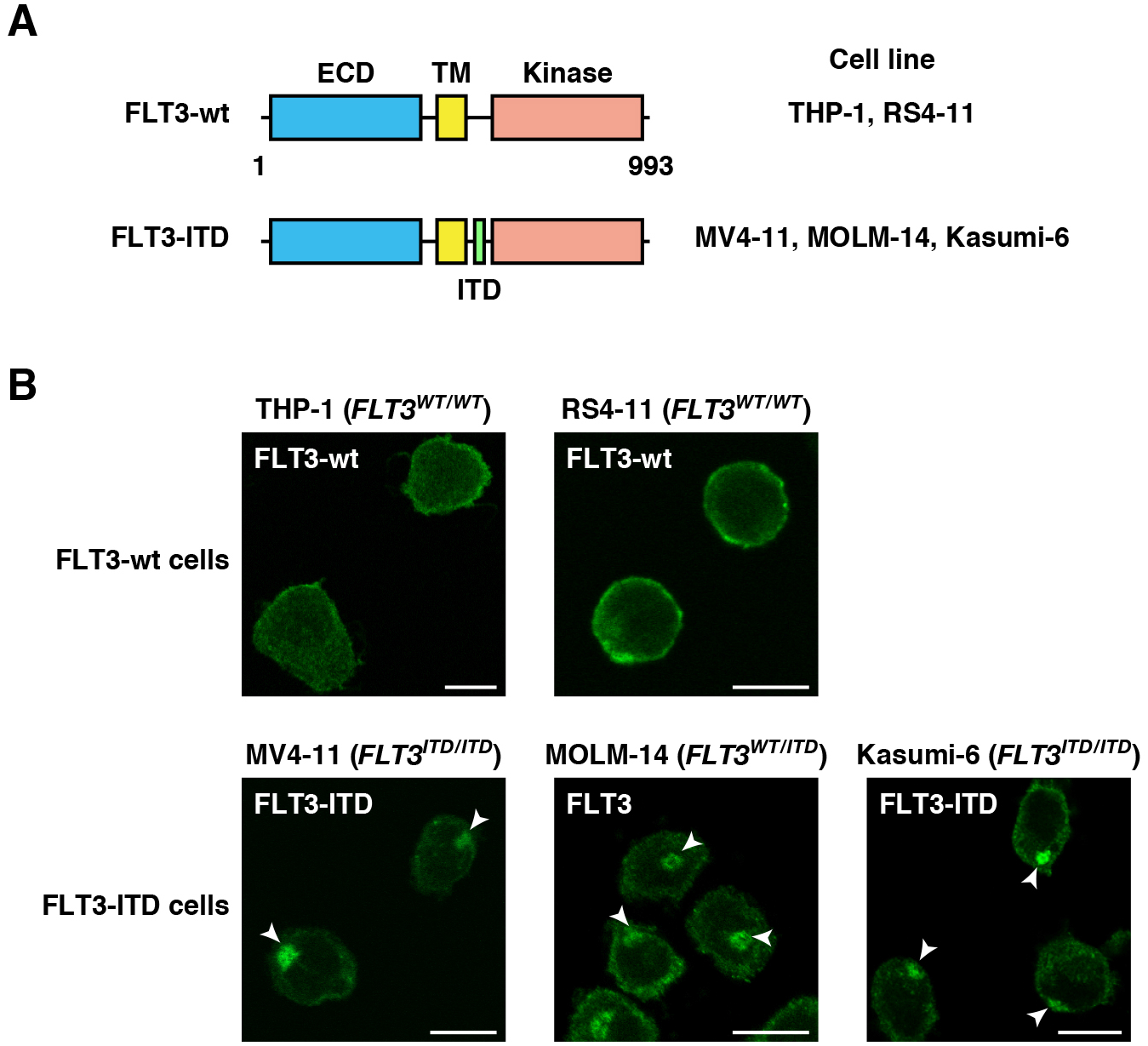
FLT3-ITD mislocalizes to the perinuclear region in AML cells. **(A)** Schematic representations of wild-type FLT3 (FLT3-wt) and a FLT3 internal tandem duplication (FLT3-ITD) mutant showing the extracellular domain (ECD, blue), the transmembrane domain (TM. yellow), the kinase domain (pink), and the ITD (green). (B) Fixed THP-L RS4-11. MV4-11. MOLM-14. or Kasumi-6 cells were permeabilized and subsequently immunostained with anti-FLT3 ECD antibody. Arrowheads indicate the perinuclear region. Bars. 10 μm. Note that FLT3-wt localized to the plasma membrane, whereas FLT3-ITD accumulated in the perinuclear region.

Recently, we reported that KIT, a type III RTK, accumulates in intracellular compartments, such as endosomal/lysosomal membrane and the Golgi apparatus, in mast cell leukemia (MCL), gastrointestinal stromal tumor (GIST), and AML (Obata et al., 2014; Obata et al., 2017; Obata et al., 2019; Saito et al., 2020). Mutant KIT in leukemia localizes to endosome-lysosome compartments through endocytosis, whereas that in GIST stops in the Golgi region during early secretory trafficking. We further showed that blockade of KIT trafficking to the signal platform inhibits oncogenic signals (Hara et al., 2017; Obata et al., 2018; Obata et al., 2019), suggesting that trafficking suppression is a novel strategy for suppression of tyrosine phosphorylation signals.

In this study, we show that endogenous FLT3-ITD aberrantly accumulates in the perinuclear region in AML cells. In our co-staining assays, we found that the perinuclear region was consistent with the Golgi region but not with the ER, endosomes, or lysosomes. The Golgi retention of FLT3-ITD is decreased by PKC412, a tyrosine kinase inhibitor (TKI), suggesting that the mutant stays in the Golgi region in a manner that is dependent on its kinase activity. Interestingly, FLT3-ITD can activate AKT and ERK in the Golgi region before reaching the PM. Inhibiting the biosynthetic trafficking of FLT3-ITD from the ER to the Golgi by brefeldin A (BFA) or 2-methylcoprophilinamide (M-COPA) can block the activation of AKT and ERK by FLT3-ITD. We also confirmed that STAT5 is activated by FLT3-ITD in the ER. Our findings provide evidence that FLT3-ITD signaling occurs on intracellular compartments, such as the Golgi apparatus and ER, in AML cells.

## Materials & Methods

### Cell culture

RS4-11, MV4-11, THP-1 (American Type Culture Collection, Manassas, VA), and MOLM-14 (Leibniz Institute DSMZ-German Collection of Microorganisms and Cell Cultures GmbH, Braunschweig, Germany) were cultured at 37°C in RPMI1640 medium supplemented with 10% fetal calf serum (FCS), penicillin/streptomycin, glutamine (Pen/Strep/Gln), and 50 μM 2-mercaptoethanol (2-ME). Kasumi-6 cells (Japanese Collection of Research Bioresources Cell Bank, Osaka, Japan) were cultured at 37°C in RPMI1640 medium supplemented with 20% FCS, 2 ng/mL granulocyte-macrophage colony-stimulating factor (Peprotech, Rocky Hill, NJ), Pen/Strept/Gln, and 50 μM 2-ME. All human cell lines were authenticated by Short Tandem Repeat analysis and tested for *Mycoplasma* contamination with a MycoAlert Mycoplasma Detection Kit (Lonza, Basel, Switzerland).

### Chemicals

PKC412 (Selleck, Houston, TX) was dissolved in dimethyl sulfoxide (DMSO). BFA (Sigma-Aldrich, St. Louis, MO) and monensin (Biomol, Hamburg, Germany) were dissolved in ethanol or methanol, respectively. M-COPA (also known as AMF-26) was synthesized as previously described (Shiina et al., 2013; Shiina et al., 2018) and dissolved in DMSO.

### Antibodies

The sources of purchased antibodies were as follows: FLT3 (S-18) and STAT5 (C-17) from Santa Cruz Biotechnology (Dallas, TX); FLT3 (8F2), FLT3[pY842] (10A8), FLT3[pY591] (54H1), AKT (40D4), AKT[pT308] (C31E5E), STAT5 (D2O6Y), STAT5[pY694] (D47E7), ERK1/2 (137F5) and ERK[pT202/pY204] (E10) from Cell Signaling Technology (Danvers, MA); TfR (ab84036), TGN46 (ab76282) and GM130 (EP892Y) from Abcam (Cambridge, UK); Calnexin (ADI-SPA-860) from Enzo (Farmingdale, NY); LAMP1 (L1418) from Sigma (St. Louis, MO) and FLT3 (MAB812) from R&D Systems (Minneapolis, MN). Horseradish peroxidase-labeled (HRP-labeled) anti-mouse IgG and anti-rabbit IgG secondary antibodies were purchased from The Jackson Laboratory (Bar Harbor, MA). Alexa Fluor-conjugated (AF-conjugated) secondary antibodies were obtained from Thermo Fisher Scientific (Rockford, IL). The list of antibodies with sources and conditions of immunoblotting and immunofluorescence is shown in Supplementary Table 1.

### Immunofluorescence confocal microscopy

Leukemia cells in suspension culture were fixed with 4% paraformaldehyde (PFA) for 20 min at room temperature, then cyto-centrifuged onto coverslips. Fixed cells were permeabilized and blocked for 30 min in phosphate-buffered saline (PBS) supplemented with 0.1% saponin and 3% bovine serum albumin (BSA), and then incubated with a primary and a secondary antibody for 1 hour each. AF647-conjugated *lectin-Helix pomatia* agglutinin (lectin-HPA, Thermo Fisher Scientific) was used for Golgi staining. After washing with PBS, cells were mounted with Fluoromount (DiagnosticBioSystems, Pleasanton, CA). For staining the extracellular domain of FLT3, living MOLM-14 cells were stained with anti-FLT3 (SF1.340) and AF488-conjugated anti-mouse IgG in PBS supplemented with 3% BSA and 0.1% sodium azide (NaN3) at 4°C for 1 hour each. Stained cells were fixed with 4% PFA for 20 min at room temperature. Confocal images were obtained with an Fluoview FV10i (Olympus, Tokyo, Japan) or a TCS SP5 II/SP8 (Leica, Wetzlar, Germany) laser scanning microscope. Composite figures were prepared with an FV1000 Viewer (Olympus), LAS X (Leica), Photoshop, and Illustrator software (Adobe, San Jose, CA).

### Western blotting

Lysates prepared in SDS-PAGE sample buffer were subjected to SDS-PAGE and electro-transferred onto PVDF membranes. Basically, 5% skimmed milk in tris-buffered saline with Tween 20 (TBS-T) was used for diluting antibodies. For immunoblotting with anti-FLT3[pY842] (10A8) or anti-FLT3[pY591], the antibody was diluted with 3% BSA in TBS-T. Immunodetection was performed with Enhanced Chemiluminescence Prime (PerkinElmer, Waltham, MA). Sequential re-probing of membranes was performed after the complete removal of primary and secondary antibodies in stripping buffer (Thermo Fisher Scientific), or inactivation of HRP by 0.1% NaN3. Results were analyzed with an LAS-3000 with Science Lab software (Fujifilm, Tokyo, Japan) or a ChemiDoc XRC+ with Image Lab software (BIORAD, Hercules, CA).

### Immunoprecipitation

Lysates from ~5 x 10^6^ cells were prepared in NP-40 lysis buffer (50 mM HEPES, pH 7.4, 10% glycerol, 1% NP-40, 4 mM EDTA, 100 mM NaF, 1 mM Na_3_VO_4_, cOmplete™ protease inhibitor cocktail (Sigma), and 1 mM PMSF). Immunoprecipitation was performed at 4°C for 3 hours using protein G Sepharose pre-coated with anti-FLT3 (S-18). Immunoprecipitates were dissolved in SDS-PAGE sample buffer.

### Cell proliferation assay

Cells were cultured with PKC412 for 48 hours. Cell proliferation was quantified using the CellTiter-GLO Luminescent Cell Viability Assay (Promega, Madison, WI), according to the manufacturer’s instructions. ATP production was measured by ARVO X3 2030 Multilabel Reader (PerkinElmer, Waltham, MA).

### Analysis of protein glycosylation

Following the manufacturer’s instructions (New England Biolabs, Ipswich, MA), NP-40 cell lysates were treated with endoglycosidases for 1 hour at 37°C. Since the FLT3 expression level in THP-1 was low for this assay, FLT3 was concentrated by immunoprecipitation with anti-FLT3 (S-18), and then treated with endoglycosidases. The reactions were stopped with SDS-PAGE sample buffer, and products were resolved by SDS-PAGE and immunoblotted.

## Results

### In human leukemia cells, wild-type FLT3 localizes to the PM, whereas FLT3-ITD accumulates in the perinuclear region

To examine the localization of endogenous FLT3, we performed confocal immunofluorescence microscopic analyses on human leukemia cell lines with an anti-FLT3 luminal-faced N-terminal region antibody. For immunostaining, we chemically fixed and permeabilized THP-1 (acute monocytic leukemia, *FLT3^WT/WT^*), RS4-11 (AML, *FLT3^WT/WT^*), MV4-11 (AML, *FLT3^ITD/ITD^*), MOLM-14 (AML, *FLT3^WT/ITD^*), and Kasumi-6 (AML, *FLT3^ITD/ITD^*) (Quentmeier et al., 2003; Furukawa et al., 2007; Wang et al., 2018; van Alphen et al., 2020; Figure 1A). In FLT3-wt leukemia cell lines (THP-1 and RS4-11), the wild-type receptor was mainly found at the PM (Figure 1B, upper panels). In sharp contrast, in *ITD*-harboring AML cells (MV4-11, MOLM-14, and Kasumi-6), the anti-FLT3 particularly stained the perinuclear region (Figure 1B, lower panels, arrowheads). The anti-FLT3 cytoplasmic domain antibody also stained the perinuclear region in these *ITD*-positive cells (Supplementary Figure S1), supporting the results of FLT3-ITD mislocalization. Since these three cell lines have different *ITD* sequences (Quentmeier et al., 2003; Furukawa et al., 2007; van Alphen et al., 2020), the accumulation of FLT3 in the perinuclear region was independent of inserted amino acid sequences but dependent on an *ITD* insertion. These results suggest that *ITD* causes FLT3 retention in the perinuclear compartment in AML cells.

### FLT3-ITD but not FLT3-wt localizes to the perinuclear Golgi region in AML cells

Next, we investigated the perinuclear region, where FLT3-ITD is found, by examining AML cell lines using co-staining assays. First, we immunostained for FLT3 (green) in conjunction with *trans-Golgi* network protein 46 kDa (TGN46, Golgi marker, red), Golgi matrix protein 130 kDa (GM130, Golgi marker, red), lectin-HPA (Golgi marker, blue), calnexin (ER marker, red), transferrin receptor (TfR, endosome marker, red), or lysosome-associated membrane protein 1 (LAMP1, lysosome marker, red) in MOLM-14 cells. As shown in Figure 2A, perinuclear FLT3 (green) was co-localized with the Golgi markers but not with the ER marker calnexin. Furthermore, localization of endosomal/lysosomal vesicles was inconsistent with that of perinuclear FLT3 (Figure 2A), indicating that FLT3-ITD localizes to the Golgi region in MOLM-14 cells. Similar results were obtained from immunofluorescence assays using both MV4-11 and Kasumi-6 cells (Figure 2B; Supplementary Figure S2A,B). In these cells, a fraction of FLT3 was found outside the ER region (Figure 2A; Supplementary Figure S2A,B), indicating that the receptor could localize in PM of *ITD*-bearing cells. We were unable to find co-localization of FLT3-wt with Golgi markers, such as lectin-HPA and GM130, in RS4-11 cells (Figure 2C; Supplementary Figure S2C), indicating that *ITD* leads FLT3 to mislocalization to the Golgi region in AML cells.

**Figure 2.**
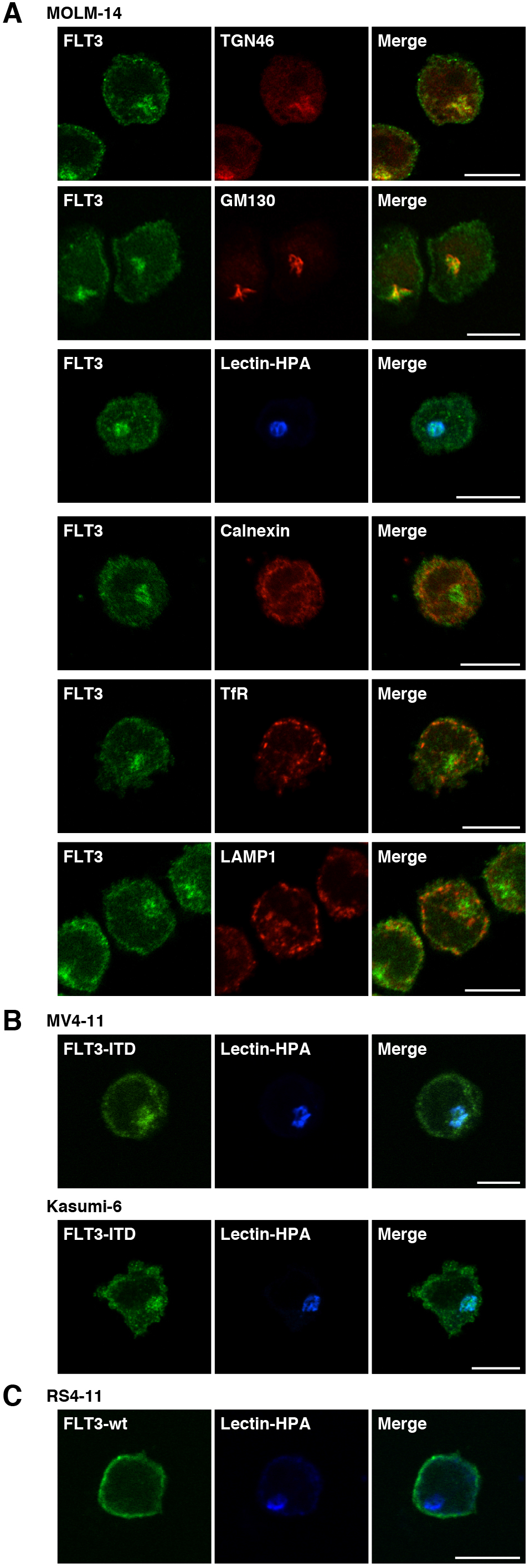
FLT3-ITD localizes to the perinuclear Golgi region in AML cells. **(A-C)** MOLM-14 **(A).** MV4-11. Kasumi-6 **(B).** or RS4-11 cells **(C)** were stained for FLT3 (green) in conjunction with the indicated organelle markers (red or blue). TGN46 (Golgi marker, red), GM130 (Golgi marker, red); lectin-HPA (Golgi marker, blue); calnexin (ER marker, red); TfR (endosome marker, red); LAMP1 (lysosome marker, red). Bars. 10 μm. Note that FLT3-ITD but not FLT3-wt accumulated in the Golgi region in AML cells.

### FLT3-ITD remains at the Golgi region in a manner dependent on its tyrosine kinase activity in AML cells

Recently, we reported that constitutively active KIT mutants in MCL, GIST, or AML accumulate in organelles in a manner dependent on their tyrosine kinase activity (Obata et al., 2014; Obata et al., 2017; Obata et al., 2019). Thus, we asked whether FLT3-ITD tyrosine kinase activity was required for retention of the mutant in the Golgi region. To answer this, we treated AML cells with midostaurin, a small molecule TKI (hereafter, referred to as PKC412), which blocks the activation of FLT3 (Furukawa et al., 2007; Pratz et al., 2010; Breitenbuecher et al., 2009; Gallogly et al., 2017; Daver et al., 2019; Kiyoi et al., 2020). Treatment of MOLM-14 cells with PKC412 suppressed the autophosphorylation of FLT3 at Tyr842 (pFLT3^Y842^) within 4 hours, resulting in a decrease in phosphor-AKT (pAKT), pERK, and pSTAT5 (Figure 3A). Treatment of MV4-11/Kasumi-6 with the TKI gave similar results (Supplementary Figure S3A,B), confirming that PKC412 suppresses the tyrosine kinase activity of FLT3-ITD and that the activation of AKT, ERK, and STAT5 is dependent on the mutant activity. As shown in Figure 3B, PKC412 suppressed the proliferation of MOLM-14. Similar to KIT (Xiang et al., 2007; Bougherara et al., 2009; Obata et al., 2014; Obata et al., 2017), upper and lower bands of FLT3 were complex-glycosylated or in a high-mannose form, respectively, since only the lower band of FLT3 was digested by endoglycosidase H treatment (Figure 3C; Supplementary Figure S3C).

**Figure 3.**
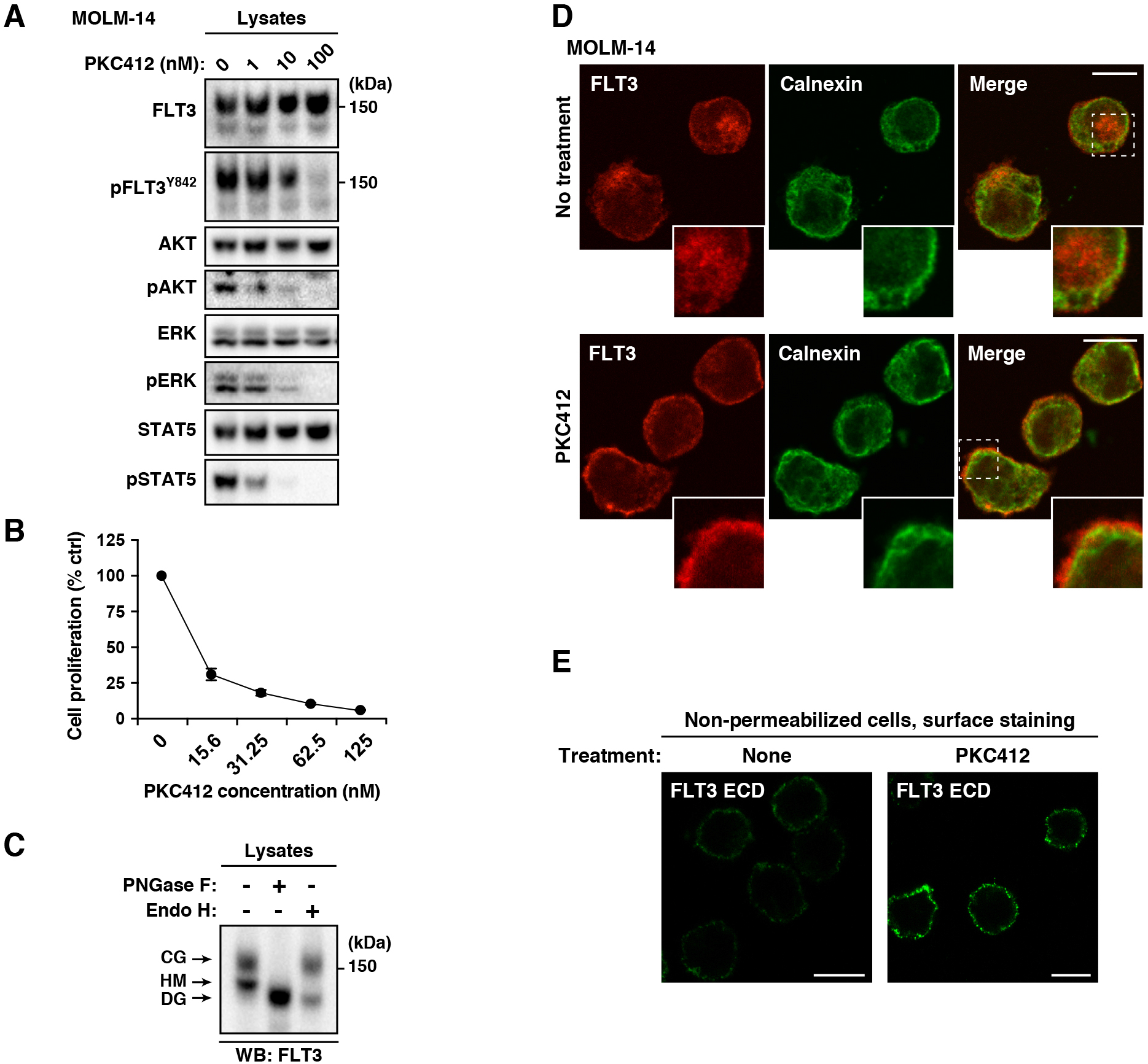
FLT3-ITD retention in the Golgi region is dependent on its tyrosine kinase activity. **(A)** MOLM-14 cells were treated for 4 hours with PKC412 (FLT3 tyrosine kinase inhibitor). Lysates were immunoblotted for FLT3. phospho-FLT3 Tyr842 (pFLT3^Y842^). AKT. pAKT. ERK. pERK. STAT5. and pSTAT5. (B) MOLM-14 cells were treated with PKC412 for 48 hours. Cell proliferation was assessed by ATP production. Results are means ± *s.d*. (n = 3). (C) Lysates from MOLM-14 were treated with peptide N-glycosidase F (PNGase F) or endoglycosidase H (endo H) then immunoblotted with anti-FLT3 antibody. CG. complex-glycosylated form; HM. high mannose form; DG. deglycosylated form. **(D.E)** MOLM-14 cells were treated with 100 nM PKC412 for 8 hours **(D)** or 16 hours **(E). (D)** Fixed cells were permeabilized. then immunostained with anti-FLT3 (red) and anti-calnexin (ER marker, green). Insets show the magnified images of the boxed area. Bars. 10 μm. **(E)** Non-permeabilized cells were immunostained with an anti-FLT3 extracellular domain (ECD) antibody. Bars. 10 μm. Note that PKC412 inactivated FLT3. then released the receptor from the Golgi region for localization to the PM.

To check the effect of PKC412 on FLT3 localization, we immunostained permeabilized MOLM-14 cells with an anti-FLT3 antibody. Interestingly, PKC412 treatment markedly diminished the FLT3 level in the Golgi region (Figure 3D, lower panels). Conversely, we found that the treatment increased the level of FLT3, probably within the PM region (Figure 3D, insets of lower panels). Previous reports showed that a kinase-dead mutation of FLT3-ITD or TKIs (AC220/crenolanib) enhance PM distribution of the mutant receptors (Schmidt-Arras et al., 2005; Jetani et al., 2018; Reiter et al., 2018; Kellner et al., 2020). Therefore, we examined the PM levels of FLT3-ITD on non-permeabilized MOLM-14 cells by staining for the FLT3 extracellular domain (ECD). As shown in Figure 3E, PKC412 treatment enhanced the PM staining of FLT3, similar to previous reports on AC220- or crenolanib-treated MV4-11 (Reiter et al., 2018; Jetani et al., 2018). These results suggest that FLT3-ITD remains in the Golgi region during secretory trafficking in a manner dependent on its kinase activity and that TKIs move the receptor to the PM.

### In AML cells, FLT3-ITD can activate STAT5, ATK, and ERK in the early secretory compartments

Finally, we examined the relationship between FLT3-ITD localization and growth signals. To determine whether FLT3-ITD activated downstream molecules, we treated AML cells for with monensin, which suppresses secretory trafficking thorough blocking Golgi export (Griffiths et al., 1983; Lievens et al., 2004; Xiang et al., 2007; Obata et al., 2014; Obata et al., 2017). As shown in Figure 4A, our immunofluorescence assay showed that FLT3 distribution other than in the Golgi region was markedly decreased in MOLM-14 cells in the presence of 100 nM monensin for 8 h, confirming the expectation that the treatment blocks Golgi export of FLT3-ITD. Next, we performed immunoblotting. Only the high-mannose form of the FLT3 bands was found in the presence of monensin (Figure 4B, top panel). In the presence of monensin, pFLT3^Y842^ was decreased (Figure 4B), whereas pFLT3^Y591^ remained (Figure 4B, bottom panel), suggesting that the treatment does not block all tyrosine phosphorylations in FLT3-ITD and that these tyrosine residues in FLT3 are regulated differently. Blocking the PM localization of FLT3-ITD caused it to be partially decreased in pAKT and pERK, but these phosphorylations and pSTAT5 remained in MOLM-14, MV4-11, and Kasumi-6 cells (Figure 4B-D). We found that FLT3 signals occurred in the presence of monensin for 24 hours (Supplementary Figure S4). These results indicate that FLT3-ITD can activate downstream before it reaches the PM.

**Figure 4.**
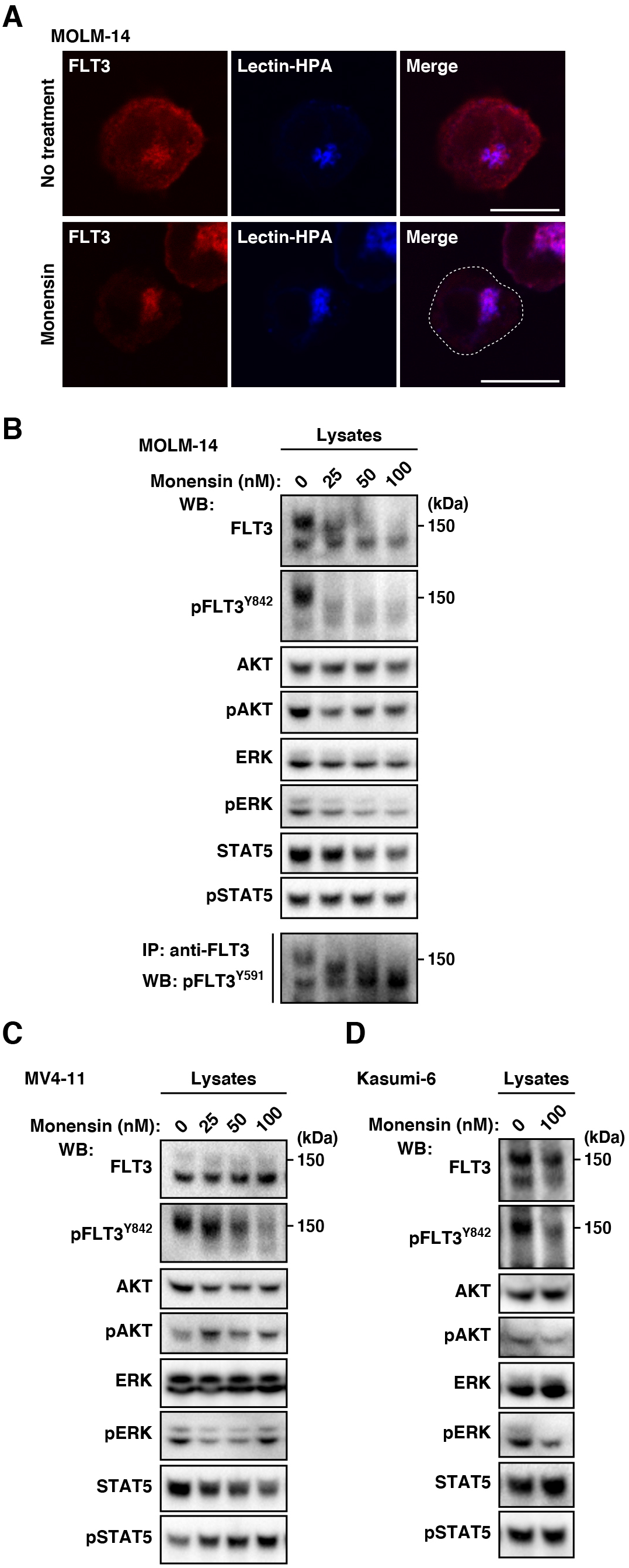
In AML cells, FLT3-ITD can activate AKT, ERK, and STAT5 before it reaches the PM. **(A.B)** MOLM-14 cells were treated with monensin (inhibitor of Golgi export) for 8 hours. (A) Cells treated with 100 nM monensin were stained with anti-FLT3 (red) and lectin-HPA (Golgi marker, blue). Dashed line, cell border. Bars. 10 μm. **(B)** Lysates were immunoblotted with the indicated antibody. To examine phospho-FLT3 Tyr59l (pFLT3^Y591^). FLT3 was immunoprecipitated, then immunoblotted. **(C.D) MV4-**11 **(C)** or Kasumi-6 cells **(D)** were treated with monensin for 8 hours, then immunoblotted. Note that FLT3-ITD in the early secretory pathway can activate downstream in AML cells.

A previous study showed that BFA, an inhibitor of ER export (Lippincott-Schwartz et al., 1989), suppresses the activation of AKT and ERK but not STAT5 in MV4-11 cells (Choudhary et al., 2009; Moloney et al., 2017). Recently, we reported that in addition to BFA, M-COPA blocks trafficking of KIT mutants from the ER (Hara et al., 2017; Obata et al., 2018; Obata et al., 2019). Thus, we treated AML cells with M-COPA as well as BFA to confirm the effect of blockade of ER export on FLT3 signaling. Our immunofluorescence assay on MOLM-14 cells showed that BFA/M-COPA treatment decreased FLT3 levels in the Golgi region within 8 hours and greatly increased the co-localization of calnexin (ER marker) with FLT3 (Figure 5A, see inset panels), confirming that these inhibitors block protein transport from the ER to the Golgi apparatus. Consistent with a previous report (Choudhary et al., 2009), pFLT3^Y842^ was diminished by BFA/M-COPA treatment, indicating that the phosphorylation does not occur in the ER. On the other hand, pFLT3^Y591^ was maintained in the ER (Figure 5B, left, bottom panel). In MV4-11 and Kasumi-6 as well as MOLM-14, FLT3-ITD in the ER was unable to activate AKT and ERK (Figure 5B-D). Blockade of ER export, however, did not inhibit STAT5 activation through FLT3-ITD (Figure 5B-D). Taken together, these results suggest that in AML cells, FLT3-ITD can activate STAT5 and AKT/ERK on the ER and the Golgi apparatus, respectively.

**Figure 5.**
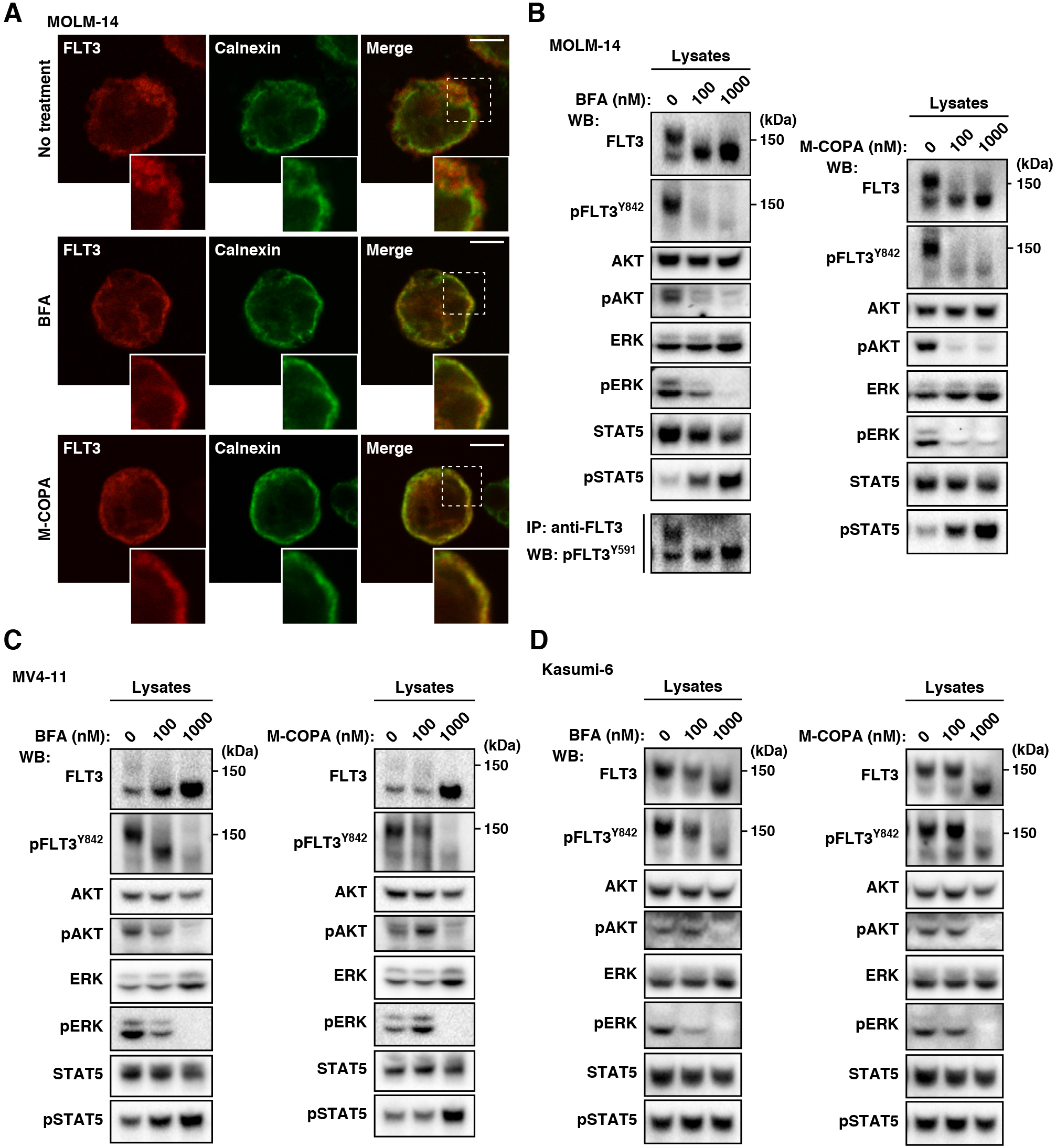
In AML cells, FLT3-ITD in the ER can activate STAT5 but not AKT or ERK. **(A-D)** MOLM-14 **(A.B).** MV4-11 **(C).** or Kasumi-6 cells **(D)** were treated with inhibitors of ER export (BFA or M-COPA) for 8 hours. **(A)** MOLM-14 cells treated with 1 μM BFA (middle panels) or 1 μM M-COPA (bottom panels) were stained with anti-FLT3 (red) and calnexin (ER marker, green). Insets show the magnified images of the boxed area. Bars. 10 μm. **(B)** Lysates were immunoblotted with the indicated antibody. To examine phospho-FLT3 Tyr59l (pFLT3^Y591^). FLT3 was immunoprecipitated, then immunoblotted. (C.D) MV4-11 (C) or Kasumi-6 cells (D) were treated with BFA or M-COPA for 8 hours, then immunoblotted. Note that BFA and M-COPA inhibited the activation of AKT and ERK but not that of STAT5 through blocking FLT3-ITD trafficking from the ER to the Golgi apparatus.

## Discussion

In this study, we demonstrate that unlike FLT3-wt (Figure 6, left), FLT3-ITD accumulates in the early secretory organelle, such as the Golgi apparatus, and in that location, causes tyrosine phosphorylation signaling in AML cells (Figure 6, right). The Golgi retention of FLT3-ITD is dependent on the tyrosine kinase activity of the mutant. TKI increases PM levels of FLT3-ITD by releasing the mutant from the Golgi apparatus. FLT3-ITD in the Golgi region can activate AKT and ERK, whereas that in the ER triggers STAT5 phosphorylation, leading to autonomous cell proliferation.

**Figure 6.**
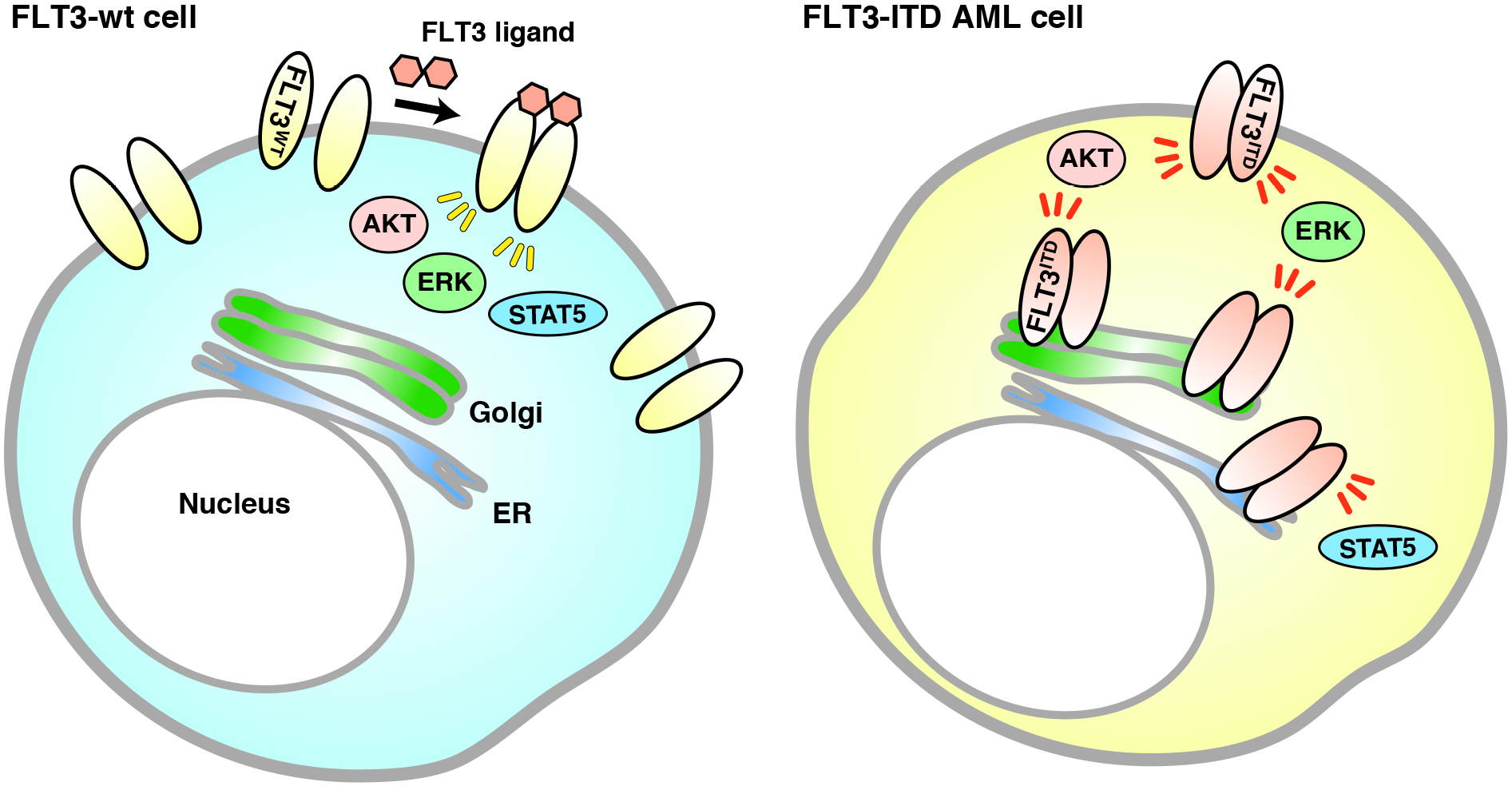
Model of FLT3-ITD signaling on intracellular compartments in AML cells. **(Left)** FLT3-wt normally moves to the PM along the secretory pathway for binding its ligand. Upon stimulation with FLT3 ligand at the cell surface, the wild-type receptor activates downstream molecules. **(Right)** FLT3-ITD is retained in the Golgi apparatus in AML cells. The mutant can activate downstream, such as AKT and ERK. in the perinuclear Golgi region but not in the ER before reaching the PM. On the other hand. FLT3-ITD activates STAT5 in the ER. where it is newly synthesized.

Recently, we reported that in MCL, KIT^D816V^ (human) or KIT^D814Y^ (mouse) activates STAT5 and AKT on the ER and endolysosomes, respectively (Obata et al., 2014; Hara et al., 2017), whereas KIT^V560G^ in MCL activates downstream at the Golgi apparatus (Obata et al., 2019). Furthermore, KIT mutants including KIT^D816V^ in cells other than MCL, such as GIST and leukemia cells, cause oncogenic signals on the Golgi apparatus (Xiang et al., 2007; Obata et al., 2017; Obata et al., 2019). As previously described (Schmidt-Arras et al., 2009; Choudhary et al., 2009; Tsitsipatis et al., 2017), we confirmed that after biosynthesis in the ER, FLT3-ITD causes STAT5 tyrosine phosphorylation in a manner similar to KIT^D816V^ in MCL. On the other hand, activation of AKT and ERK through FLT3-ITD is similar to activation through the KIT mutant in GIST in that it occurs on the Golgi apparatus. A recent report showed that FLT3^D835Y^ is also found in endomembranes (Rudorf et al., 2019). As described above, since the signal platform for kinases may be affected by its mutation site, there is great interest in carrying out a further investigation to determine whether FLT3^D835Y^ causes growth signaling on the ER, Golgi, or endosome/lysosomes.

Recently, novel protein interactions and downstream molecules for FLT3 which support cancer cell proliferation have been identified. An FLT3 mutant is found to activate Rho kinase through activation of RhoA small GTPase, resulting in myeloproliferative disease development (Mali et al., 2011). GADS physically associates with FLT3-ITD, and the interaction enhances downstream activation (Chougule et al., 2016). Analysis of spatio-temporal associations of FLT3 mutants with these functional interactors is an attractive possibility.

Previous studies reported that other RTK mutants, such as FGFR3^K650E^ in multiple myeloma, RET multiple endocrine neoplasia type 2B (RET^MEN2B^), and PDGFRA^Y289C^, are also tyrosine-phosphorylated via the secretory pathway (Ronchetti et al., 2001; Lievens et al., 2004; Lievens et al., 2006; Gibbs & Legeai-Mallet, 2007; Runeberg-Roos et al., 2007; Toffalini & Demoulin, 2010; Ip et al., 2018; Schmidt-Arras & Böhmer, 2020). Signal transduction from the secretory compartments may be a characteristic feature of a large number of RTK mutants. Early secretory compartments can be subdivided into ER, the ER-Golgi intermediate compartment, *cis-, medial-Golgi* cisternae, and TGN, and others. It would be interesting to identify the sub-compartment at which RTK mutants are retained for precise understanding of the mechanism of growth signaling. Three-dimensional super-resolution confocal microscopic analysis on cancer cells is now underway.

Golgi retention of FLT3-ITD is dependent on receptor tyrosine kinase activity. As with previous reports (Reiter et al., 2018; Kellner et al., 2020), we confirmed that a TKI increased PM localization of FLT3-ITD, indicating that these inhibitors can release the mutant from the Golgi region for localization to the PM. Previous reports together with the results of our studies showed that TKIs increase the PM levels of RTK mutants, such as EGFR^T790M^, KIT^D816V^, and PDGFRA^V561G^ (Bougherara et al., 2013; Obata et al., 2014; Watanuki et al., 2014; Bahlawane et al., 2015; Obata et al., 2019). Enhancement of PM distribution with TKIs may be a common feature of RTK mutants. Furthermore, recent studies showed that the effect of chimeric antigen receptor T-cell therapy and antigen-dependent cell cytotoxicity using anti-FLT3 is enhanced by increasing the PM levels of FLT3-ITD through TKI treatment (Durben et al., 2015; Jetani et al., 2018; Reiter et al., 2018; Wang et al., 2018). Combining TKIs together with immunotherapy will lead to improvements in the prognosis of cancer patients.

TKIs and antibodies against RTKs have been developed for suppression of growth signals in cancers. In this study, blockade of the ER export of FLT3-ITD with BFA/M-COPA greatly decreased tyrosine phosphorylation signals in AML cells. Since the bioavailability of M-COPA *in vivo* is higher than that of BFA and can be orally administered to animals, we will investigate the anti-cancer effect of the compound on AML-bearing mice. Together with the results of previous reports (Joffre et al., 2011; Williams et al., 2012; Larrue et al., 2015; Hara et al., 2017; Obata et al., 2018; Zappa et al., 2018; Obata et al., 2019; Prieto-Dominguez et al., 2019), our findings suggest that an intracellular trafficking blockade of RTK mutants could be a third strategy for inhibition of oncogenic signaling.

In conclusion, we show that in AML cells, the perinuclear region where FLT3-ITD accumulates is the Golgi apparatus. Similar to KIT mutants in GISTs, FLT3-ITD is retained at the Golgi region in a manner dependent on its kinase activity, but TKI releases the mutant to the PM. Our findings provide new insights into the role of FLT3-ITD in autonomous AML cell growth. Moreover, from a clinical point of view, our findings offer a new strategy for AML treatment through blocking the involvement of FLT3-ITD in secretory trafficking.

## Abbreviations

AML: acute myeloid leukemia
BFA: brefeldin A
EGFR: epidermal growth factor receptor
ER: endoplasmic reticulum
ERK: extracellular signal-regulated kinase
FGFR: fibroblast growth factor receptor
FLT3: fms-like tyrosine kinase 3
GADS: GRB2-related adaptor downstream of Shc
GIST: gastrointestinal stromal tumor
ITD: internal tandem duplication
JM: juxta-membrane
MCL: mast cell leukemia
M-COPA: 2-methylcoprophilinamide
PDGFR: platelet-derived growth factor receptor
pFLT3: phospho-FLT3
PM: plasma membrane
RET: rearranged during transfection
RTK: receptor tyrosine kinase
STAT: signal transducers and activators of transcription
TGN: *trans*-Golgi network
TKI: tyrosine kinase inhibitor
wt: wild-type.

## Acknowledgments

The authors thank Dr. Yusuke Furukawa, Dr. Jiro Kikuchi (Jikei Medical University) and Dr. Mitsutoshi Tsukimoto (Tokyo University of Science) for their helpful advice and sharing of materials. This work was supported by a grant-in-aid for Scientific Research from the Japan Society for the Promotion of Science (18K07208 to YO, 19H03722 to TN, and 20K08719 to RA), by research grants from the Kawano Masanori Memorial Public Interest Incorporated Foundation for Promotion of Pediatrics, the Friends of Leukemia Research Fund, and the Ichiro Kanehara Foundation for the Promotion of Medical Sciences and Medical Care (to YO).

## Authors’ contributions

KY performed and analyzed the data from all experiments and wrote the manuscript. IS supervised the total synthesis of M-COPA and edited the manuscript. TM, ST, and YM carried out the synthesis of M-COPA and helped to draft the manuscript. MN performed immunoblotting and edited the manuscript. MS, KO, and TS provided advice on the design of the *in vitro* experiments. TN provided advice on the design of the *in vitro* experiments and edited the manuscript. RA conceived and supervised the project, analyzed the data and wrote the manuscript. YO conceived, designed, performed and analyzed the data from all experiments, and wrote the manuscript. All authors read and approved the final version.

## Ethics approval and consent to participate

Not applicable.

## Consent for publication

Not applicable.

## Competing interests

The authors declare that they have no competing interests.

## Notes

### Competing Interest Statement

The authors have declared no competing interest.

## References

Bahlawane C, Eulenfeld R, Wiesinger MY, Wang J, Muller A, Girod A, Nazarov PV, Felsch K, Vallar L, Sauter T, et al. Constitutive activation of oncogenic PDGFRα-mutant proteins occurring in GIST patients induces receptor mislocalisation and alters PDGFRα signalling characteristics. Cell Commun. Signal. 2015; 13: doi: 10.1186/s12964-015-0096-8.

Bougherara H, Subra F, Crépin R, Tauc P, Auclair C, Poul MA. The aberrant localization of oncogenic kit tyrosine kinase receptor mutants is reversed on specific inhibitory treatment. Mol. Cancer Res. 2009; 7: 1525–1533. doi: 10.1158/1541-7786.MCR-09-0138.

Bougherara H, Georgin-Lavialle S, Damaj G, Launay JM, Lhermitte L, Auclair C, Arock M, Dubreuil P, Hermine O, Poul MA. Relocalization of KIT D816V to cell surface after dasatinib treatment: potential clinical implications. Clin. Lymphoma Myeloma Leuk. 2013; 13: 62–69. doi: 10.1016/j.clml.2012.08.004.

Breitenbuecher F, Markova B, Kasper S, Carius B, Stauder T, Böhmer FD, Masson K, Rönnstrand L, Huber C, Kindler T, et al. A novel molecular mechanism of primary resistance to FLT3-kinase inhibitors in AML. Blood. 2009; 113: 4063–4073. doi: 10.1182/blood-2007-11-126664.

Chatterjee A, Ghosh J, Ramdas B, Mali RS, Martin H, Kobayashi M, Vemula S, Canela VH, Waskow ER, Visconte V, et al. Regulation of stat5 by FAK and PAK1 in oncogenic FLT3-and KIT-driven leukemogenesis. Cell Rep. 2014; 9: 1333–1348. doi: 10.1016/j.celrep.2014.10.039.

Choudhary C, Olsen JV, Brandts C, Cox J, Reddy PN, Böhmer FD, Gerke V, Schmidt-Arras DE, Berdel WE, Müller-Tidow C, et al. Mislocalized activation of oncogenic RTKs switches downstream signaling outcomes. Mol. Cell. 2009; 36: 326–339. doi: 10.1016/j.molcel.2009.09.019.

Chougule RA, Cordero E, Moharram SA, Pietras K, Rönnstrand L, Kazi JU. Expression of GADS enhances FLT3-induced mitogenic signaling. Oncotarget. 2016; 7: 14112–14124. doi: 10.18632/oncotarget.7415.

Daver N, Schlenk RF, Russell NH, Levis MJ. Targeting *FLT3* mutations in AML: review of current knowledge and evidence. Leukemia. 2019; 33: 299–312. doi: 10.1038/s41375-018-0357-9.

Durben M, Schmiedel D, Hofmann M, Vogt F, Nübling T, Pyz E, Bühring HJ, Rammensee HG, Salih HR, Große-Hovest L, et al. Characterization of a bispecific FLT3 X CD3 antibody in an improved, recombinant format for the treatment of leukemia. Mol. Ther. 2015; 23: 648–655. doi: 10.1038/mt.2015.2.

Furukawa Y, Vu HA, Akutsu M, Odgerel T, Izumi T, Tsunoda S, Matsuo Y, Kirito K, Sato Y, Mano H, et al. Divergent cytotoxic effects of PKC412 in combination with conventional antileukemic agents in FLT3 mutation-positive versus -negative leukemia cell lines. Leukemia. 2007; 21: 1005–1014. doi: 10.1038/sj.leu.2404593.

Gallogly MM, Lazarus HM, Cooper BW. Midostaurin: a novel therapeutic agent for patients with FLT3-mutated acute myeloid leukemia and systemic mastocytosis. Ther. Adv. Hematol. 2017; 8: 245–261. doi: 10.1177/2040620717721459.

Gibbs L, Legeai-Maller L. FGFR3 intracellular mutations induce tyrosine phosphorylation in the Golgi and defective glycosylation. Biochim. Biophys. Acta. 2007; 1773: 502–512. doi: 10.1016/j.bbamcr.2006.12.010.

Griffiths G, Quinn P, Warren G. Dissection of the Golgi complex. I. Monensin inhibits the transport of viral membrane proteins from *medial* to *trans* Golgi cisternae in baby hamster kidney cells infected with Semliki Forest virus. J. Cell Biol. 1983; 96: 835–850. doi: 10.1083/jcb.96.3.835.

Hara Y, Obata Y. Horikawa K, Tasaki Y, Suzuki K, Murata T, Shiina I, Abe R. M-COPA suppresses endolysosomal Kit-Akt oncogenic signalling through inhibiting the secretory pathway in neoplastic mast cells. PLOS ONE. 2017; 12: doi: 10.1371/journal.pone.0175514.

Heydt Q, Larrue C, Saland E, Bertoli S, Sarry JE, Besson A, Manenti S, Joffre C, Mansat-De Mas V. Oncogenic FLT3-ITD supports autophagy via ATF4 in acute myeloid leukemia. Oncogene. 2018; 37: 787–797. doi: 10.1038/onc.2017.376.

Ip CKM, Ng PKS, Jeong KJ, Shao SH, Ju Z, Leonard PG, Hua X, Vellano CP, Woessner R, Sahni N, et al. Neomorphic *PDGFRA* extracellular domain driver mutations are resistant to PDGFRA targeted therapies. Nat. Commun. 2018; 9: 4583. doi: 10.1038/s41467-018-06949-w.

Jetani H, Garcia-Cadenas I, Nerreter T, Thomas S, Rydzek J, Meijide JB, Bonig H, Herr W, Sierra J, Einsele H, et al. CAR T-cells targeting FLT3 have potent activity against FLT3 ^-^ ITD ^+^ AML and act synergistically with the FLT3-inhibitor crenolanib. Leukemia. 2018; 32: 1168–1179. doi: 10.1038/s41375-018-0009-0.

Joffre C, Barrow R, Ménard L, Calleja V, Hart IR, Kermorgant S. A direct role for Met endocytosis in tumorigenesis. Nat. Cell Biol. 2011; 13: 827–837. doi: 10.1038/ncb2257.

Kazi JU, Rönnstrand L. FMS-like tyrosine kinase 3/FLT3: from basic science to clinical implications. Physiol. Rev. 2019; 99: 1433–1466. doi: 10.1152/physrev.00029.2018.

Kellner F, Keil A, Schindler K, Tschongov T, Hünninger K, Loercher H, Rhein P, Böhmer SA, Böhmer FD, Müller JP. Wild-type FLT3 and FLT3 ITD exhibit similar ligand-induced internalization characteristics. J. Cell. Mol. Med. 2020; 24: 4668–4676. doi: 10.1111/jcmm.15132.

Kiyoi H, Ohno R, Ueda R, Saito H, Naoe T. Mechanism of constitutive activation of FLT3 with internal tandem duplication in the juxtamembrane domain. Oncogene. 2002; 21: 2555–2563. doi: 10.1038/sj.onc.1205332.

Kiyoi H, Kawashima N, Ishikawa Y. *FLT3* mutations in acute myeloid leukemia: Therapeutic paradigm beyond inhibitor development. Cancer Sci. 2020; 111: 312–322. doi: 10.1111/cas.14274.

Koch S, Jacobi A, Ryser M, Ehninger G, Thiede C. Abnormal localization and accumulation of FLT3-ITD, a mutant receptor tyrosine kinase involved in leukemogenesis. Cells Tissues Organs. 2008; 188: 225–235. doi: 10.1159/000118788.

Köthe S, Müller JP, Böhmer SA, Tschongov T, Fricke M, Koch S, Thiede C, Requardt RP, Rubio I, Böhmer FD. Features of Ras activation by a mislocalized oncogenic tyrosine kinase: FLT3 ITD signals through K-Ras at the plasma membrane of acute myeloid leukemia cells. J. Cell Sci. 2013; 126: 4746–4755. doi: 10.1242/jcs.131789.

Larrue C, Saland E, Vergez F, Serhan N, Delabesse E, Mansat-De Mas V, Hospital MA, Tamburini J, Manenti S, Sarry JE, et al. Antileukemic activity of 2-deoxy-d-glucose through inhibition of N-linked glycosylation in acute myeloid leukemia with *FLT3-ITD* or *c-KIT* mutations. Mol. Cancer Ther. 2015; 14: 2364–2373. doi: 10.1158/1535-7163.MCT-15-0163.

Lemmon MA, Schlessinger J. Cell signaling by receptor tyrosine kinases. Cell. 2010; 141: 1117–1134. doi: 10.1016/j.cell.2010.06.011.

Lievens PM, Mutinelli C, Baynes D, Liboi E. The kinase activity of fibroblast growth factor receptor 3 with activation loop mutations affects receptor trafficking and signaling. J. Biol. Chem. 2004; 279: 43254–43260. doi: 10.1074/jbc.M405247200.

Lievens PM, Roncador A, Liboi E. K644E/M FGFR3 mutants activate Erk1/2 from the endoplasmic reticulum through FRS2 alpha and PLC gamma-independent pathways. J. Mol. Biol. 2006; 357: 783–792. doi: 10.1016/j.jmb.2006.01.058.

Lippincott-Schwartz J., Yuan LC, Bonifacino JS, Klausner RD. Rapid redistribution of Golgi proteins into the ER in cells treated with brefeldin A: evidence for membrane cycling from Golgi to ER. Cell. 1989; 56: 801–813. doi: 10.1016/0092-8674(89)90685-5.

Mali RS, Ramdas B, Ma P, Shi J, Munugalavadla V, Sims E, Wei L, Vemula S, Nabinger SC, Goodwin CB, et al. Rho kinase regulates the survival and transformation of cells bearing oncogenic forms of KIT, FLT3, and BCR-ABL. Cancer Cell. 2011; 20: 357–369. doi: 10.1016/j.ccr.2011.07.016.

Meshinchi S, Appelbaum FR. Structural and functional alterations of FLT3 in acute myeloid leukemia. Clin. Cancer Res. 2009; 15: 4263–4269. doi: 10.1158/1078-0432.CCR-08-1123.

Moloney JN, Stanicka J, Cotter TG. Subcellular localization of the FLT3-ITD oncogene plays a significant role in the production of NOX-and p22^phox^-derived reactive oxygen species in acute myeloid leukemia. Leuk. Res. 2017; 52: 34–42. doi: 10.1016/j.leukres.2016.11.006.

Obata Y, Toyoshima S, Wakamatsu E, Suzuki S, Ogawa S, Esumi H, Abe R. Oncogenic Kit signals on endolysosomes and endoplasmic reticulum are essential for neoplastic mast cell proliferation. Nat. Commun. 2014; 5: 5715. doi: 10.1038/ncomms6715.

Obata Y, Horikawa K, Takahashi T, Akieda Y, Tsujimoto M, Fletcher JA, Esumi H, Nishida T, Abe R. Oncogenic signaling by Kit tyrosine kinase occurs selectively on the Golgi apparatus in gastrointestinal stromal tumors. Oncogene. 2017; 36: 3661–3672. doi: 10.1038/onc.2016.519.

Obata Y, Horikawa K, Shiina I, Takahashi T, Murata T, Tasaki Y, Suzuki K, Yonekura K, Esumi H, Nishida T, et al. Oncogenic Kit signalling on the Golgi is suppressed by blocking secretory trafficking with M-COPA in GISTs. Cancer Lett. 2018; 415: 1–10. doi: 10.1016/j.canlet.2017.11.032.

Obata Y, Hara Y, Shiina I, Murata T, Tasaki Y, Suzuki K, Ito K, Tsugawa S, Yamawaki K, Okamoto K, et al. N822K-or V560G-mutated KIT activation occurs preferentially in lipid rafts of the Golgi apparatus in leukemia cells. Cell Commun. Signal. 2019; 17: 114. doi: 10.1186/s12964-019-0426-3.

Pratz KW, Sato T, Murphy KM, Stine A, Rajkhowa T, Levis M. FLT3-mutant allelic burden and clinical status are predictive of response to FLT3 inhibitors in AML. Blood. 2010; 115: 1425–1432. doi: 10.1182/blood-2009-09-242859.

Prieto-Dominguez N, Parnell C, Teng Y. Drugging the small GTPase pathways in cancer treatment: promises and challenges. Cells. 2019; 8: 255. doi: 10.3390/cells8030255.

Quentmeier H, Reinhardt J, Zaborski M, Drexler HG. FLT3 mutations in acute myeloid leukemia cell lines. Leukemia. 2003, 17, 120–124. doi: 10.1038/sj.leu.2402740.

Reiter K, Polzer H, Krupka C, Maiser A, Vick B, Rothenberg-Thurley M, Metzeler KH, Dörfel D, Salih HR, Jung G, et al. Tyrosine kinase inhibition increases the cell surface localization of FLT3-ITD and enhances FLT3-directed immunotherapy of acute myeloid leukemia. Leukemia. 2018; 32: 313–322. doi: 10.1038/leu.2017.257.

Ronchetti D, Greco A, Compasso S, Colombo G, Dell’Era P, Otsuki T, Lombardi L, Neri A. Deregulated FGFR3 mutants in multiple myeloma cell lines with t(4;14): comparative analysis of Y373C, K650E and the novel G384D mutations. Oncogene. 2001; 20: 3553–3562. doi: 10.1038/sj.onc.1204465.

Rudorf A, Müller TA, Klingeberg C, Kreutmair S, Poggio T, Gorantla SP, Rückert T, Schmitt-Graeff A, Gengenbacher A, Paschka P, et al. NPM1c alters FLT3-D835Y localization and signaling in acute myeloid leukemia. Blood. 2019; 134: 383–388. doi: 10.1182/blood.2018883140.

Runeberg-Roos P, Virtanen H, Saarma M. RET(MEN 2B) is active in the endoplasmic reticulum before reaching the cell surface. Oncogene. 2007; 26: 7909–7915. doi: 10.1038/sj.onc.1210591.

Saito Y, Takahashi T, Obata Y, Nishida T, Ohkubo S, Nakagawa F, Serada S, Fujimoto M, Ohkawara T, Nishigaki T, et al. TAS-116 inhibits oncogenic KIT signalling on the Golgi in both imatinib-naïve and imatinib-resistant gastrointestinal stromal tumours. Br. J. Cancer. 2020; 122: 658–667. doi: 10.1038/s41416-019-0688-y.

Schmidt-Arras DE, Böhmer A, Markova B, Choudhary C, Serve H, Böhmer FD. Tyrosine phosphorylation regulates maturation of receptor tyrosine kinases. Mol. Cell. Biol. 2005; 25: 3690–3703. doi: 10.1128/MCB.25.9.3690-3703.2005.

Schmidt-Arras D, Böhmer SA, Koch S, Müller JP, Blei L, Cornils H, Bauer R, Korasikha S, Thiede C, Böhmer FD. Anchoring of FLT3 in the endoplasmic reticulum alters signaling quality. Blood. 2009; 113: 3568–3576. doi: 10.1182/blood-2007-10-121426.

Schmidt-Arras D, Böhmer FD. Mislocalisation of activated receptor tyrosine kinases - Challenges for cancer therapy. Trends Mol. Med. 2020; 26: 833–847. doi: 10.1016/j.molmed.2020.06.002.

Shiina I, Umezaki Y, Ohashi Y, Yamazaki Y, Dan S, Yamori T. Total synthesis of AMF-26, an antitumor agent for inhibition of the Golgi system, targeting ADP-ribosylation factor 1. J. Med Chem. 2013; 56: 150–159. doi: 10.1021/jm301695c.

Shiina I, Umezaki Y, Murata T, Suzuki K, Tonoi T. Asymmetric total synthesis of (+)-coprophilin. Synthesis. 2018; 50: 1301–1306. 10.1055/s-0036-1591866.

Takahashi S. Mutations of FLT3 receptor affect its surface glycosylation, intracellular localization, and downstream signaling. Leuk. Res. Rep. 2019; 13: 100187. doi: 10.1016/j.lrr.2019.100187.

Toffalini F, Demoulin JB. New insights into the mechanisms of hematopoietic cell transformation by activated receptor tyrosine kinases. Blood. 2010; 116: 2429–2437. doi: 10.1182/blood-2010-04-279752.

Tsapogas P, Mooney CJ, Brown G, Rolink A. The cytokine Flt3-ligand in normal and malignant hematopoiesis. Int. J. Mol. Sci. 2017; 18: 1115. doi: 10.3390/ijms18061115.

Tsitsipatis D, Jayavelu AK, Müller JP, Bauer R, Schmidt-Arras D, Mahboobi S, Schnöder TM, Heidel F, Böhmer FD. Synergistic killing of FLT3ITD-positive AML cells by combined inhibition of tyrosine-kinase activity and N-glycosylation. Oncotarget. 2017; 8: 26613–26624. doi: 10.18632/oncotarget.15772.

van Alphen C, Cloos J, Beekhof R, Cucchi DGJ, Piersma SR, Knol JC, Henneman AA, Pham TV, van Meerloo J, Ossenkoppele GJ, et al. Phosphotyrosine-based phosphoproteomics for target identification and drug response prediction in AML cell lines. Mol. Cell. Proteomics. 2020; 19: 884–899. doi: 10.1074/mcp.RA119.001504.

Wang Y, Xu Y, Li S, Liu J, Xing Y, Xing H, Tian Z, Tang K, Rao Q, Wang M, et al. Targeting FLT3 in acute myeloid leukemia using ligand-based chimeric antigen receptor-engineered T cells. J. Hematol. Oncol. 2018; 11: 60. doi: 10.1186/s13045-018-0603-7.

Watanuki Z, Kosai H, Osanai N, Ogama N, Mochizuki M, Tamai K, Yamaguchi K, Satoh K, Fukuhara T, Maemondo M, et al. Synergistic cytotoxicity of afatinib and cetuximab against EGFR T790M involves Rab11-dependent EGFR recycling. Biochem. Biophys. Res. Commun. 2014; 455: 269–276. doi: 10.1016/j.bbrc.2014.11.003.

Williams AB, Li L, Nguyen B, Brown P, Levis M, Small D. Fluvastatin inhibits FLT3 glycosylation in human and murine cells and prolongs survival of mice with FLT3/ITD leukemia. Blood. 2012; 120: 3069–3079. doi: 10.1182/blood-2012-01-403493.

Xiang Z, Kreisel F, Cain J, Colson AL, Tomasson MH. Neoplasia driven by mutant *c-KIT* is mediated by intracellular, not plasma membrane, receptor signaling. Mol. Cell. Biol. 2007; 27: 267–282. doi: 10.1128/MCB.01153-06.

Yamamoto Y, Kiyoi H, Nakano Y, Suzuki R, Kodera Y, Miyawaki S, Asou N, Kuriyama K, Yagasaki F, Shimazaki C, et al. Activating mutation of D835 within the activation loop of FLT3 in human hematologic malignancies. Blood. 2001; 97: 2434–2439. doi: 10.1182/blood.v97.8.2434.

Zappa F, Failli M, De Matteis MA. The Golgi complex in disease and therapy. Curr. Opin. Cell Biol. 2018; 50: 102–116. doi: 10.1016/j.ceb.2018.03.005.

