## Supplemental Figures for "FLT3-ITD transduces autonomous growth signals during its biosynthetic trafficking in acute myelogenous leukemia cells"

### Supplementary Figures

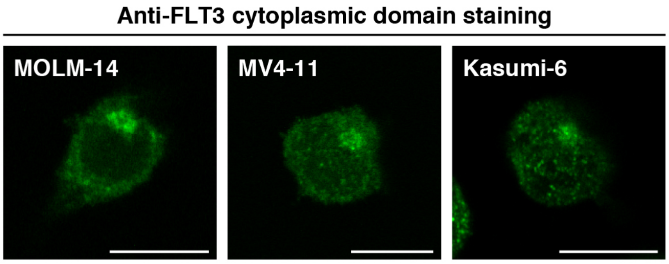


**Supplementary Figure S1. An anti-FLT3 cytoplasmic domain antibody stains the perinuclear region of MOLM-14, MV4-11, and Kasumi-6 cells.** FLT3-ITD-harboring AML cell lines were immunostained with an anti-FLT3 cytoplasmic domain antibody. Bars, 10 µm.

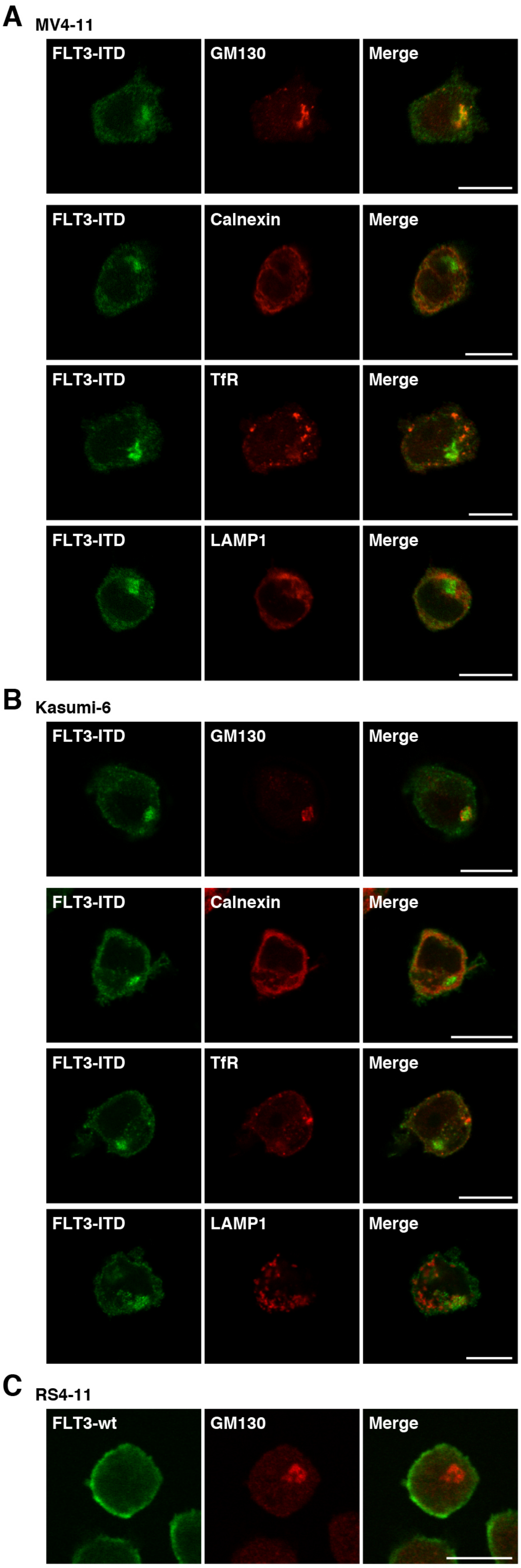


**Supplementary Figure S2. FLT3-ITD localizes in the Golgi region in MV4-11 and Kasumi-6 cells.** (**A-C**) MV4-11 (**A**), Kasumi-6 (**B**), or RS4-11 cells (**C**) were immunostained for FLT3 (green) in conjunction with the indicated organelle markers (red). GM130 (Golgi marker); calnexin (ER marker); TfR (endosome marker); LAMP1 (lysosome marker). Bars, 10 μm.

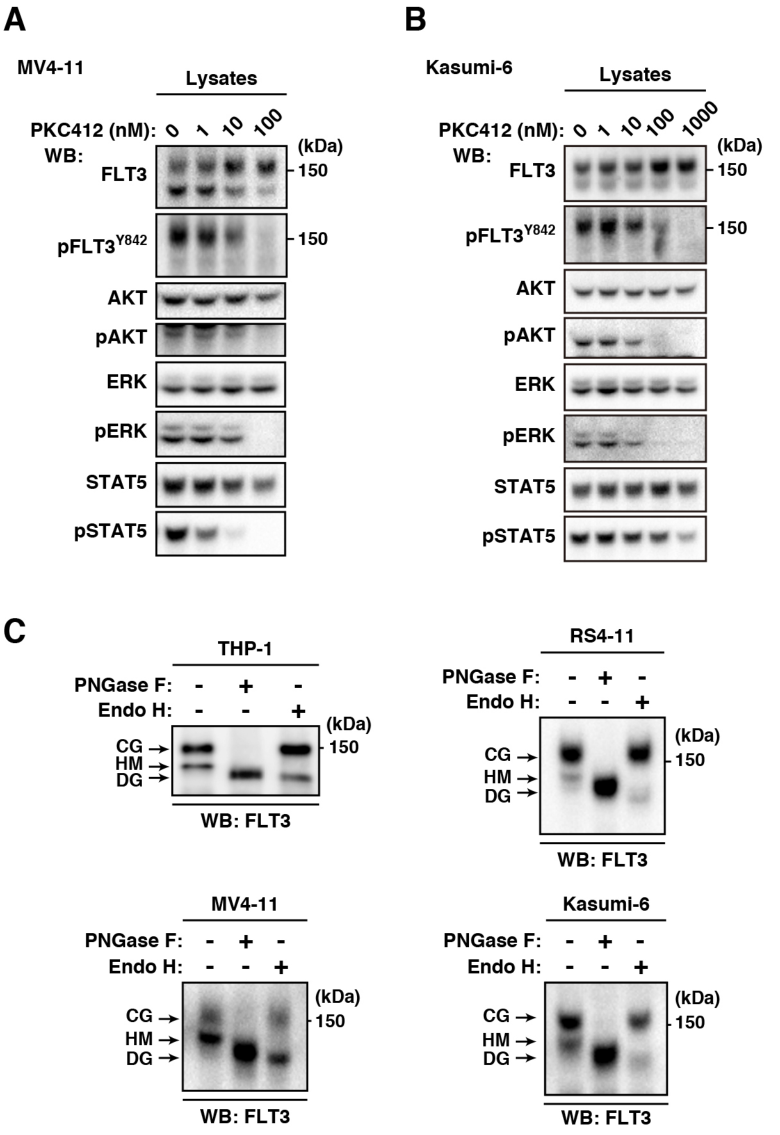


**Supplementary Figure S3. PKC412 inhibits the phosphorylation of AKT, ERK, and STAT5 thorough suppressing FLT3-ITD activation in MV4-11 and Kasumi-6 cells.** (**A**,**B**) MV4-11 (**A**) or Kasumi-6 cells (**B**) were treated for 4 hours with PKC412 (inhibitor of FLT3 tyrosine kinase). Lysates were immunoblotted for FLT3, phospho-FLT3 Tyr842 (pFLT3$^{Y842}$), AKT, pAKT, ERK, pERK, STAT5, and pSTAT5. (**C**) Lysates from leukemia cell lines were treated with peptide N-glycosidase F (PNGase F) or endoglycosidase H (endo H) then immunoblotted with anti-FLT3. CG, complex-glycosylated form; HM, high mannose form; DG, deglycosylated form.

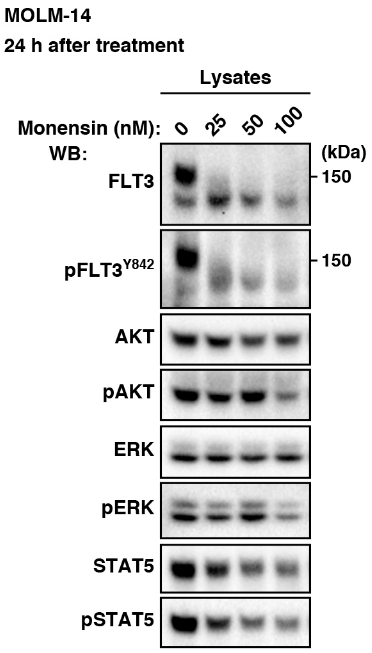


**Supplementary Figure S4. In AML cells, FLT3-ITD can activate AKT, ERK, and STAT5 before it reaches the PM.** MOLM-14 cells were treated with monensin (inhibitor of Golgi export) for 24 hours. Lysates were immunoblotted for FLT3, phospho-FLT3 Tyr842 (pFLT3$^{Y842}$), AKT, pAKT, ERK, pERK, STAT5, and pSTAT5.

### Supplementary Table 1. List of antibodies

| Antibody | Clone/catalog # | Distribution source | WB | IF |
|---|---|---|---|---|
| AKT | 40D4 | Cell Signaling Technology | 1/1000 | - |
| AKT [pT308] | C31E5E | Cell Signaling Technology | 1/1000 | - |
| Calnexin | ADI-SPA-860 | Enzo | - | 1/250 |
| FLT3 | S-18 | Santa Cruz Biotechnology | 1/1000 | - |
| FLT3 | 8F2 | Cell Signaling Technology | 1/1000 | 1/250 |
| FLT3 | MAB812 | R&D Systems | - | 1/250 |
| FLT3 | SF1.340 | Santa Cruz Biotechnology | - | 1/100~1/250 |
| ERK1/2 | 137F5 | Cell Signaling Technology | 1/1000~1/2000 | - |
| ERK [pT202/pY204] | E10 | Cell Signaling Technology | 1/1000 | - |
| FLT3 [pY842] | 10A8 | Cell Signaling Technology | 1/1000 | - |
| FLT3 [pY591] | 54H1 | Cell Signaling Technology | 1/1000 | - |
| GM130 | EP892Y | Abcam | - | 1/250 |
| LAMP1 | L1418 | Sigma-Aldrich | - | 1/250 |
| STAT5 | C-17 | Santa Cruz Biotechnology | 1/1000 | - |
| STAT5 | D2O6Y | Cell Signaling Technology | 1/1000 | - |
| STAT5 [pY694] | D47E7 | Cell Signaling Technology | 1/1000 | - |
| TfR | ab84036 | Abcam | - | 1/250 |
| TGN46 | ab76282 | Abcam | - | 1/100 |
| HRP donkey anti-mouse IgG | 715-035-151 | Jackson Laboratory | 1/2000 | - |
| HRP donkey anti-rabbit IgG | 711-035-152 | Jackson Laboratory | 1/2000 | - |
| AF488 donkey anti-mouse IgG | A21202 | Thermo Fisher Scientific | - | 1/250 |
| AF488 donkey anti-rabbit IgG | A21206 | Thermo Fisher Scientific | - | 1/250 |
| AF568 donkey anti-mouse IgG | A10037 | Thermo Fisher Scientific | - | 1/250 |
| AF568 donkey anti-rabbit IgG | A10042 | Thermo Fisher Scientific | - | 1/250 |
| AF647 lectin-HPA | L32454 | Thermo Fisher Scientific | - | 1/100~1/250 |

**Supplementary Table 1 |** List of antibodies. The list shows antibodies with sources and conditions of Western blotting (WB) and immunofluorescence (IF).
